# Female manipulation of offspring sex ratio in the gregarious parasitoid *Anaphes flavipes* from a new two-generation approach

**DOI:** 10.1101/2021.02.22.432331

**Authors:** Alena Samková, Jan Raška, Jiří Hadrava, Jiří Skuhrovec, Petr Janšta

## Abstract

Both theoretical and empirical work suggests that offspring sex ratio has important consequences on fitness. Within insects, gregarious parasitoids with haplodiploid sex determination represent an ideal model for studying the decision-making process behind the assignment of offspring sex. To gain insight into the offspring sex ratio of gregarious parasitoids, we performed experiments on *Anaphes flavipes*, interpreting our results through a two-generation approach. We confirm the existence of a relationship between offspring sex ratio and clutch size: the proportion of males increases with larger clutch size. Based on this finding, we assumed that the proportion of males among one female’s offspring would also increase with external factors such as a low population density of the host or the presence of the host’s predator, which may pressure the mothers to lay a higher-sized clutch. Contrary to our initial expectations, we show that if it is the pressure of external factors that leads to an increase in clutch size, these larger clutches tend to be more female-biased and the overall offspring sex ratio of a particular female does not change. While in our previous work, we showed that higher clutch sizes reduce body sizes of the offspring and their future fertility, here we conclude that the differences in fertility affect the offspring sex ratio. Taken together, we highlight our two-generation approach which reveals that while the above external factors do not affect the sex ratio of *A. flavipes* in the F1 generation, they do have an effect in the F2 generation.

## INTRODUCTION

Gregarious parasitoids with more tolerant (Schmidt & Smith 1986) and/or less mobile (Boivin & Barren 2000) larvae developing together in one host provide a unique opportunity for studies of the evolution of reproductive behaviour (Rosenheim 1996). Upon finding and accepting a host, parasitoid females face an important decision about how many offspring they will lay into one host (Klomp & Teerink 1967, Charnov et al. 1981). Moreover, females of parasitoids with haplodiploid sex determination (Cook 1993, Godfray 1994) can manipulate not only the clutch size but also the offspring sex ratio by controlling the fertilisation of their eggs (King 1989). For gregarious parasitoids, some combinations of the number and sex ratio of offspring developing in a single host are more advantageous than other (Klomp & Teerink 1967, Waage & Lane 1984) and therefore favoured by natural selection to maximize individual fitness (Waage & Ming 1984). Compared to solitary parasitoids with random mating and an unbiased offspring sex ratio, the sex ratio of gregarious parasitoids is usually female-biased, with mating taking place on the host before dispersion (Hamilton 1967, Forsse et al. 1992, Godfray 1994). As a result, gregarious parasitoids will tend to balance an advantageous reproductive strategy of obtaining as many fertilized female offspring with the least possible number of male offspring, just enough for them to able to fertilize all the female offspring in the clutch (Hardy 1992).

The sex ratio of gregarious parasitoids is, among other factors, often influenced by the host (Pandey & Singh 1999, Harvey et al. 2000, Vacari et al. 2012), maternal age (Pandey & Singh 1998, Rungrojwanich & Walter 2000) or size (Carbone et al. 2007, Farahani 2016), female wasp density (Antolin 1992), brood and clutch size (Vet et al. 1994, Gu et al. 2003), as well as genetic factors (Stouthamer et al. 1992, Grbić et al. 1992). The most frequently studied factor and one which apparently has a significant influence on the sex ratio of the parasitoid’s offspring is host size (e.g., Waage 1982, King 1989, Ueno 1999). Parasitoids preferably oviposit female offspring into large hosts (Charnov 1981, Henri & van Veen 2011, Mawela et al. 2003), because these usually require a higher amount of nourishment during larval development than male offspring (Godfray 1994). In gregarious parasitoids, the amount of food for larval development is determined not only by the host size, but also by the mother’s decision on the clutch size (Klomp & Teerink 1962, Godfray 1991, Koppik et al. 2014). More males and larger clutch sizes could be expected (Yu et al. 2003) when food resources are limited, because the total fitness gained from one host may still increase because of the larger clutch size, at least until a maximum is reached (Van Alphe & Visser 1990), but exceeding the nutritive capacity of the host causes the larvae to starve (Brodeur 2006). We propose to focus on examining the parasitoid sex ratio relative to clutch size, in response to a changing external environment.

The species *Anaphes flavipes* (Förster, 1841) (Hymenoptera: Mymaridae) is an idiobiont gregarious egg parasitoid with haplodiploid sex determination (Anderson & Paschke 1968). The females can lay one to seven offspring at any sex ratio into a single host, resulting in 35 possible combinations in a clutch (Anderson & Paschke 1968). By changing clutch size, the females are able to respond to different external conditions (Samková et al. 2019b) and we predict that they can also change the offspring sex ratio. Importantly, *A. flavipes* is a potential agent for biological control against major crop pests of the genera *Lema* and *Oulema* (Coleoptera: Chrysomelidae) from Europe and North America (Linnaeus, 1758) (Dysart et al. 1973, Samková et al. 2017, Skuhrovec et al. 2018).

In this study, we measured the effect of clutch size on the sex ratio of offspring developing in one host to assess the above-mentioned assumption that larger clutch sizes should contain more male offspring (Yu et al. 2003). Secondly, we examined the offspring sex ratio in individual females under the pressure of external factors that affect clutch size (different host population density and the presence of the host’s predator; Samková et al. 2019b). Again, the tested assumption was whether the proportion of male offspring will increase with larger clutch size. However, since we previously showed that these external factors help determine the body size and fertility of the F1 generation (Samková et al. 2019b), we took this one step further and examined the effect of fertility on the sex ratio, discussing the results of our experiments from two generation approach, i.e. the perspective of both F1 and F2 generations.

## MATERIAL AND METHODS

### Parasitoids

*Anaphes flavipes* were reared from host eggs (*Oulema* spp.) collected in cereal fields (barley and wheat) in two localities in Prague-Suchdol, Czech Republic (GPS: 50.1385N, 14.3695E; 50.1367N, 14.3638E) from the end of April until the end of June in the years 2012, 2015, 2016 identically as in our previous studies (Samková et al. 2019a, b, c, 2020, recorded in Suppl. materials). The parasitised host eggs were stored in Petri dishes with moistened filter paper until adult wasps emerged. These “wild” wasps were used as an initial population to rear next generations of parasitoids in the environmental chamber at 22±2°C, 40–60% relative humidity and continuous illumination. Subsequent generations of females and males were used for experiments. Mated females (not older than 24 hours post emergence) were placed in Petri dishes with host eggs. Before the start of the experiment and during the experiment, the females were not fed, and they had free access to water (modified from Samková et al. 2019a, b, c, 2020).

### Host species

The *Oulema* species complex, including two ecologically very close species of *O. duftschmidi* Redtenbacher, 1874 and *O. melanopus* Linnaeus, 1758, was used identically as in our previous studies (Samková et al. 2019a b c, 2020) and in other studies (Anderson & Paschke 1968, 1969), because these species occur together in the same localities at the same time (Van de Vijver et al. 2019, Samková et al. 2020) and are distinguished only on the basis of genital preparation (Bezděk & Baselga 2015). We consider these species particularly safe and convenient for our experiments because 1) their host eggs do not differ in length and weight (Samková et al. 2019a), 2) the wasps *A. flavipes* are not host-specific (Samková et al. 2020) and 3) the offspring of these two host species do not differ in size (Samková et al. 2019a) and 4) show no morphological abnormalities (unpublish.). In the present study, the host culture was established from adults collected at the same time as host eggs in two localities in the Czech Republic (in Prague-Suchdol [identical to those of the parasitic wasps] and Police nad Metují [GPS: 50.5277N, 16.2456E]), using a net or individual sampling. The *Oulema* individuals were kept in plastic boxes (16.7 x 17.4 x 6.4 cm) with moistened filter paper. Adults were fed grain leaves and had unlimited access to water. They were allowed to lay their eggs on cereal leaves in an environmental chamber at 22±2°C, relative humidity of 40–60% and 16:8 L:D cycle. Host eggs were removed along with a 1 cm long piece of leaf and then used in the experiments. Since *A. flavipes* females refuse host eggs older than 72 hours (Anderson & Paschke 1968), we used eggs not older than 24 hours (modified from Samková et al. 2019a, b, c, 2020).

#### Laboratory experiments

All laboratory experiments were performed in Petri dishes (diameter: 8.5 cm) in a thermal cabinet at 22±2°C, 40–60% relative humidity and under a 16:8 (L:D) photoperiod. Individual parasitized host eggs were moved into 1.5 ml plastic tubes on the 9^th^ or 10^th^ day after parasitisation and stored at the same temperature in a thermal cabinet. For each parasitised host egg, the number and sex ratio of emerged wasps were measured. Data from these experiments were also used for studies by Samková et al. (2019a, b) focusing on body size, fertility and factors influencing the clutch size of *A. flavipes* females.

### 1 Clutch size

Each female (n = 100) had 12 host eggs available for parasitisation for 8 hours in Petri dishes. The sex ratio in relation to the number of hatched offspring in each host egg was measured.

### 2 Factors affecting the clutch size

#### Host population density

Different host population densities were simulated in the laboratory using three different treatments, similarly to our previous study (Samková et al. 2019b). In the first treatment (low host population density), 9 host eggs (3 host eggs every 8 hours) were offered to each wasp (n = 37) for parasitisation during a 24-hour period. In the second treatment (medium host population density), 24 host eggs (8 host eggs every 8 hours) were offered to each wasp (n = 44) for parasitisation during a 24-hour period. In the third treatment (high host population density) 30 host eggs were offered to each wasp (n = 46) for parasitisation during a 24-hour period. The sex of all the offspring developing in each host egg was recorded.

#### The presence of a predator

The effect of predation on host eggs was simulated by the presence of an adult seven-spot ladybird, *Coccinella septempunctata* Linnaeus, 1758, as in our previous study (Samková et al. 2019b). The predator was placed in a Petri dish (8.5 cm in diameter) with a moistened filter paper but without a wasp or any host eggs for 2 hours, in order to allow it to leave chemical cues. After 2 hours, the predator was placed in a 1.5 ml Eppendorf tube closed by a mesh (100 μm mesh size), and the tube was placed back into the Petri dish. Mated females of *A. flavipes* (n = 56) and 12 eggs of *Oulema* sp. were placed for 8 hours into a Petri dish with a predator in the Eppendorf tube. The sex ratio of all hatched offspring in each host egg was recorded for each female. The control wasp group (n = 100) received 12 host eggs for parasitisation for 8 hours without a predator.

### 3 Brood size

Each female (n = 100) had 12 host eggs available for parasitisation for 8 hours in Petri dishes. The sex ratio of the total number of hatched offspring by one female were measured.

### Statistical data processing

Software R version 4.0.1 (R. Core Team 2019) was used for all statistical analyses. The sex ratio of each mother’s offspring was assessed by generalized linear models for a binomial distribution (GLM-b); and brood size (i.e. total number of offspring of each mother) was always used as an explanatory variable in these models. Offspring sex ratio in each host was assessed by generalized mixed-effect models for a binomial distribution (GLMM-b, package lme4, Bates et al. 2014), in which the identity of each mother was included as a random factor. The effects of brood size and/or clutch size (i.e. number of offspring in each host) were used as explanatory variables in modified models, adjusted for the effects of host density and the presence of a predator on offspring sex ratio. The most suitable model was selected according to the results of likelihood ratio tests (chi-square).

Additional analyses included generalized linear models for a negative binomial distribution (GLM-nb, package MASS, Ripley et al. 2013), used for clutch size as a dependent variable.

Confidence intervals for offspring sex ratio were calculated in the R package Hmisc (Harrell & Harrell 2019).

## RESULTS

### 1) Clutch size

The sex ratio of wasps hatched from a single host egg depended on the clutch size. With a larger clutch size, the percentage of males significantly increased (GLMM-b, χ^2^_(1)_ = 13.043, p < 0.001) (Fig. 1; Suppl. Mat. 1.)

**Fig. 1.**
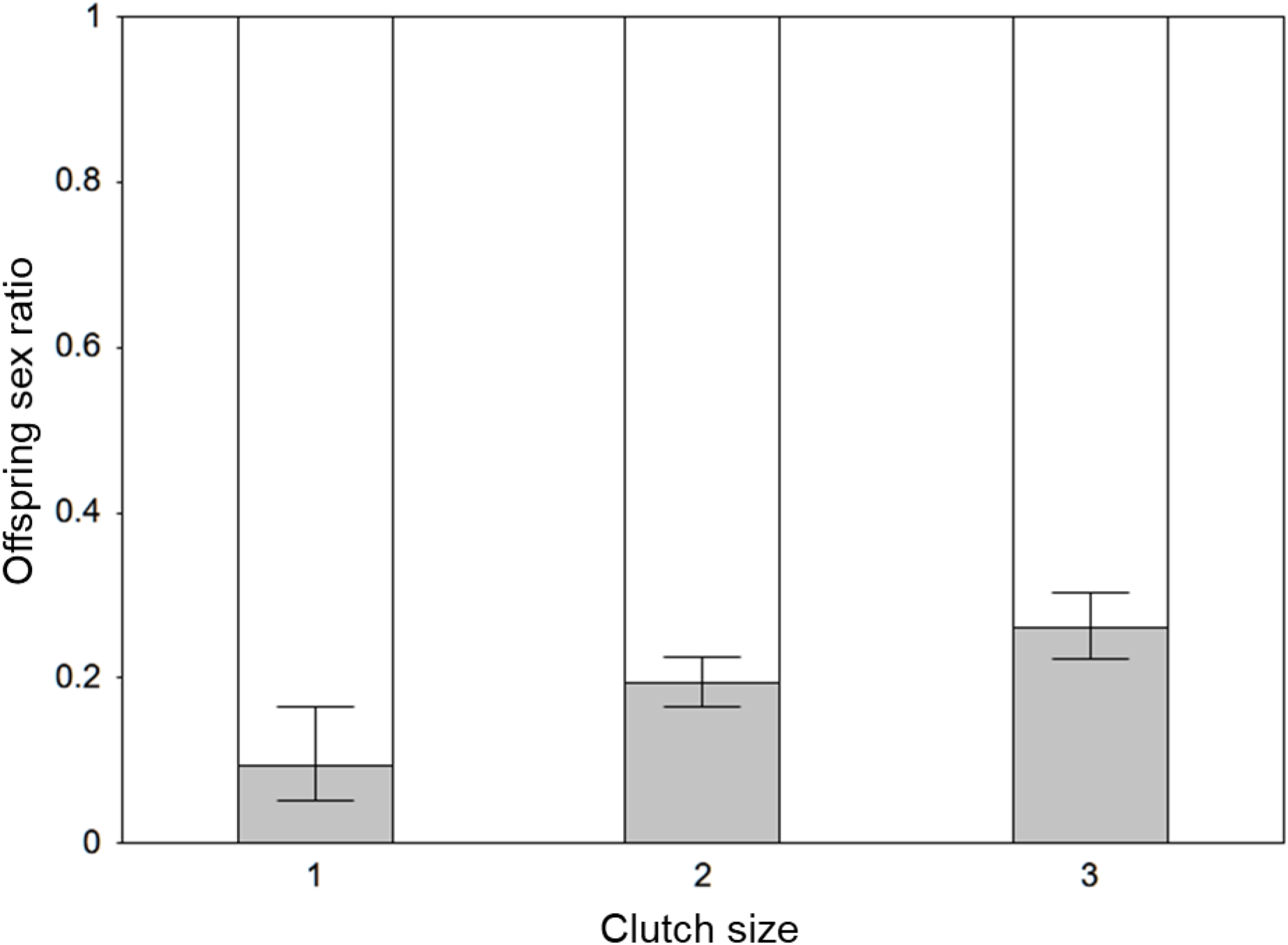
*Anaphes flavipes* offspring sex ratio (mean, 95% confidence interval) depending on clutch size (i.e. the number of offspring in a single host). Grey – males, white – females.

### 2) Factors affecting the clutch size

#### Host population density

After filtering out the effect of brood size, the sex ratio of *A. flavipes* offspring was not influenced by host population density (GLM-b, χ^2^_(2)_ = 0.362, p = 0.834; Fig. 2; Suppl. Mat. 2.). Furthermore, clutch size did influence the sex ratio of offspring in each host, but host population density did not (GLMM-b, χ^2^_(2)_ = 1.712, p = 0.425).

**Fig. 2.**
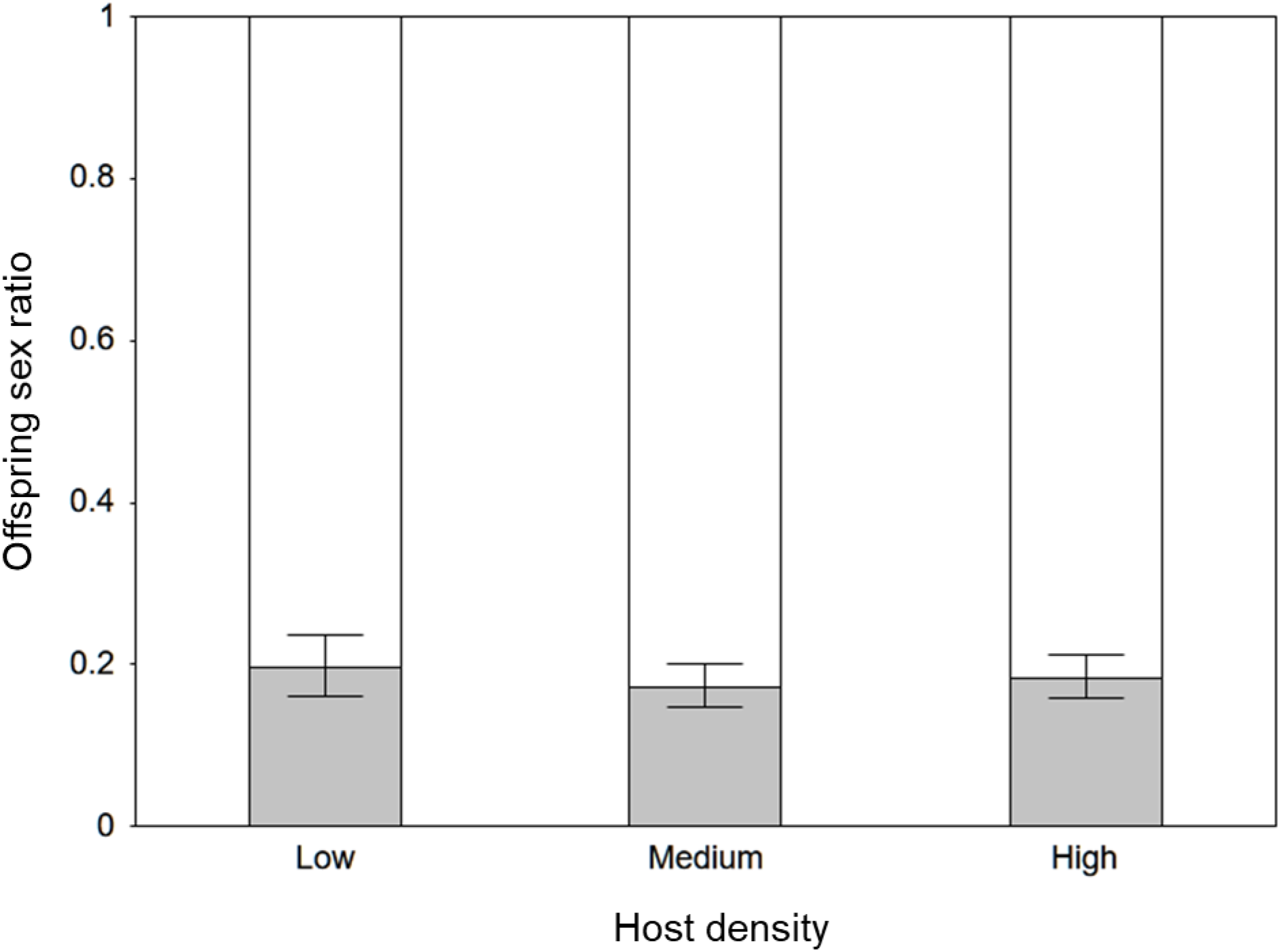
*Anaphes flavipes* offspring sex ratio (mean, 95% confidence interval) depending on the density of available hosts. Grey – males, white – females.

#### The presence of a predator

The sex ratio of *A. flavipes* offspring from one female was not significantly different between the treatment with the host egg predator present and the control without a predator (GLM-b, χ^2^_(1)_ = 1.185, p = 0.276; Fig. 3; Suppl. Mat. 3.). Additionally, the presence of a predator did not affect the sex ratio of offspring developing in one host after controlling for the effect of brood size and clutch size (GLMM-b, χ^2^_(1)_ = 0.258, p = 0.612).

**Fig. 3.**
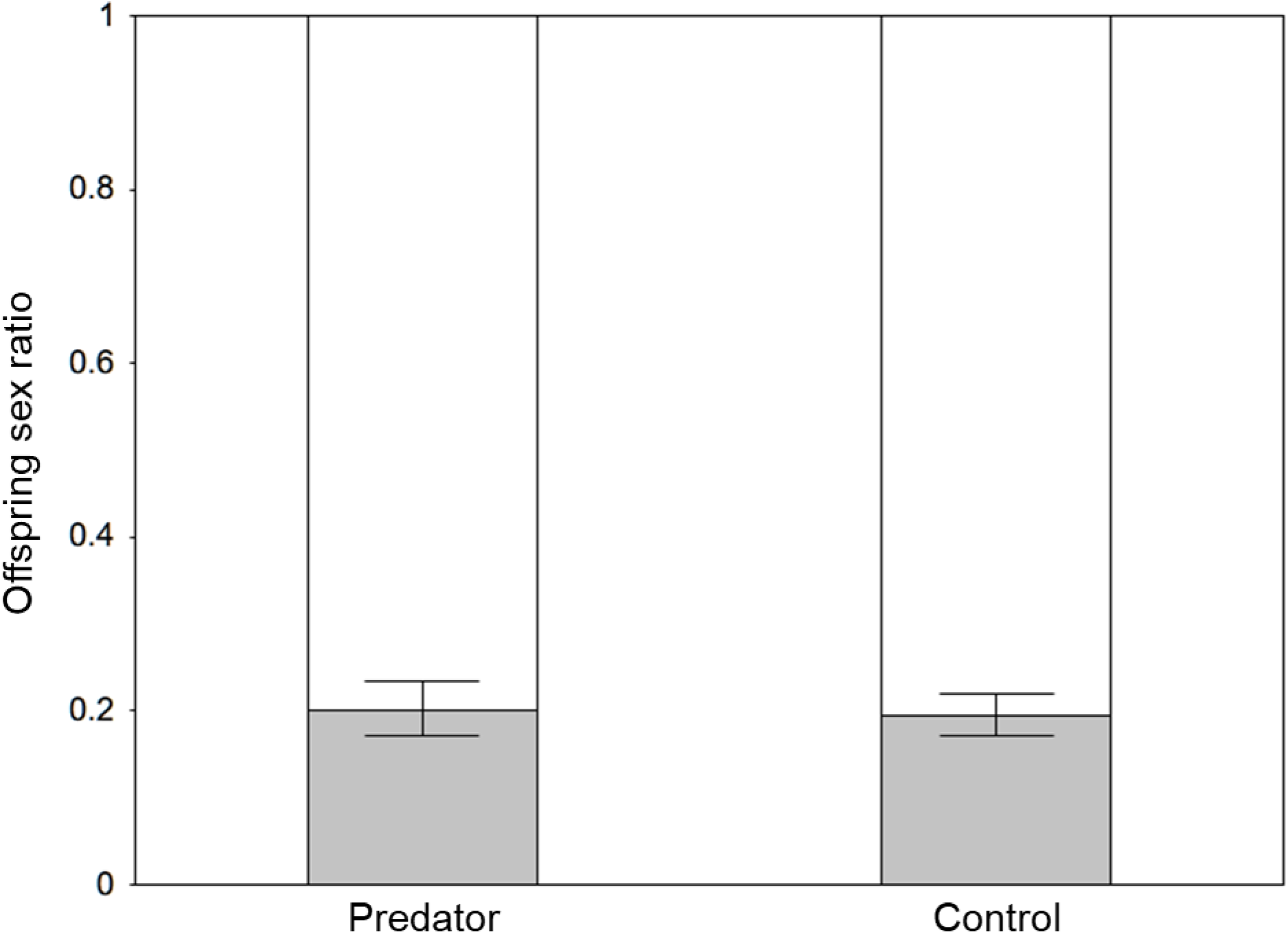
*Anaphes flavipes* offspring sex ratio (mean, 95% confidence interval) depending on the presence of a potential predator (*C. septempunctata*). Grey – males, white – females.

### 3) Fertility

The correlation between the sex ratio of the offspring of one mother and her total fertility was statistically significant: with higher brood size, the proportion of male offspring decreased (GLM-b, χ^2^_(1)_ = 9.613, p = 0.002; Fig. 4; Suppl. Mat. 4.). Females with a higher fertility also had generally larger clutches (GLM-nb, χ^2^_(1)_ = 194.8, p < 0.001), but their clutches were more female-biased than those of less fertile females (GLMM-b, χ^2^_(1)_ = 3.828, p = 0.05).

**Fig. 4.**
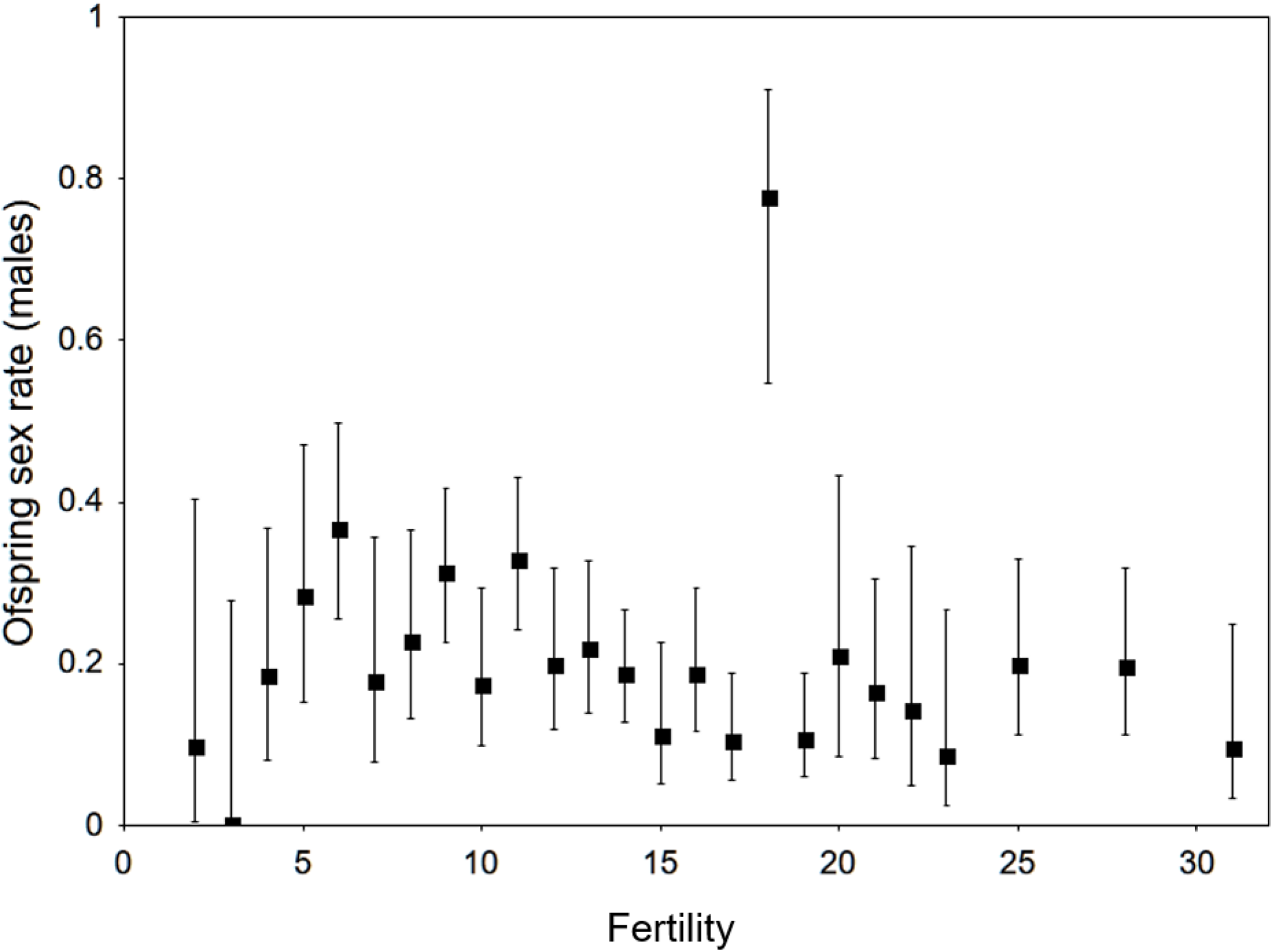
Percentage of males in *A. flavipes* offspring (mean, 95% confidence interval) depending on their fertility (i.e. the total offspring of each mother).

## DISCUSSION

Gregarious parasitoids determine the amount of food for their offspring not only by their host choice, but also by altering clutch size (Bai et al. 1992, Cloutier et al. 2000, Samková et al. 2019a). With a larger clutch size, the amount of food resources per individual offspring decreases (Godfray 1991, Koppik et al. 2014) and the number of males per clutch increases (Yu et al. 2003, this study), because male fitness is less affected by food availability during larval development than female fitness (Godfray 1994). In this study on *A. flavipes* we found that the proportion of male offspring in a clutch increased with larger clutch size, similarly to the results in the species *Habrobracon hebetor* (Say, 1836) (Yu et al. 2003). However, ‘overshooting’ male production at larger clutch sizes could actually harm the female’s interest, since females maximize their fitness by maximizing the number of mated female offspring (Hamilton 1967). Therefore, the number of males should be just enough to ensure the fertilization of all female offspring (Hardy 1992). The ovipositing females must thus find an optimal balance between the two selection pressures (Griffiths & Godfray 1988).

In this study we also examined the offspring sex ratio of *A. flavipes* under the pressure of two external factors that affect their clutch size: host population density and the presence of predator of the host (Fig. 5). Previously, we showed that lower host population density and presence of predator of host correlated with increased clutch size (Samková et al. 2019b). Although offspring sex ratio of many parasitoids such as *Pteromalus apum* (Kraft and van Nouhuys 2013), *Cotesia flavipes* (Vacari et al. 2012), or *Nasonia vitripennis* (Flanagan et al. 1998) varies based on their clutch sizes under different population densities, the sex ratio of *A. flavipes* offspring laid by one female is not altered by the different population density of the host. This is in concordance with a study by Yu et al. (2003) done on the gregarious parasitoid *Habrobracon hebetor* (Say, 1836), where the sex ratio of larger clutch sizes shifted towards males, but the sex ratio of all offspring laid by one female was not affected by host density. Likewise, when the host’s predator is present, the parasitoids developing into hosts become a potential intraguild prey (Polis et al. 1989, Nakashima & Senoo 2003). Some parasitoids react to this by recognizing the chemical marks of predators (Meisner et al. 2011) and avoiding parasitisation (Nakashima & Senoo 2003, Nakashima et al. 2006), other may lay a larger clutch size (*A. flavipes*; Samková et al. 2019b). However, here we show that concerning sex ratio, the mothers maintain the same proportion of male offspring regardless of the presence of predator.

**Fig. 5.**
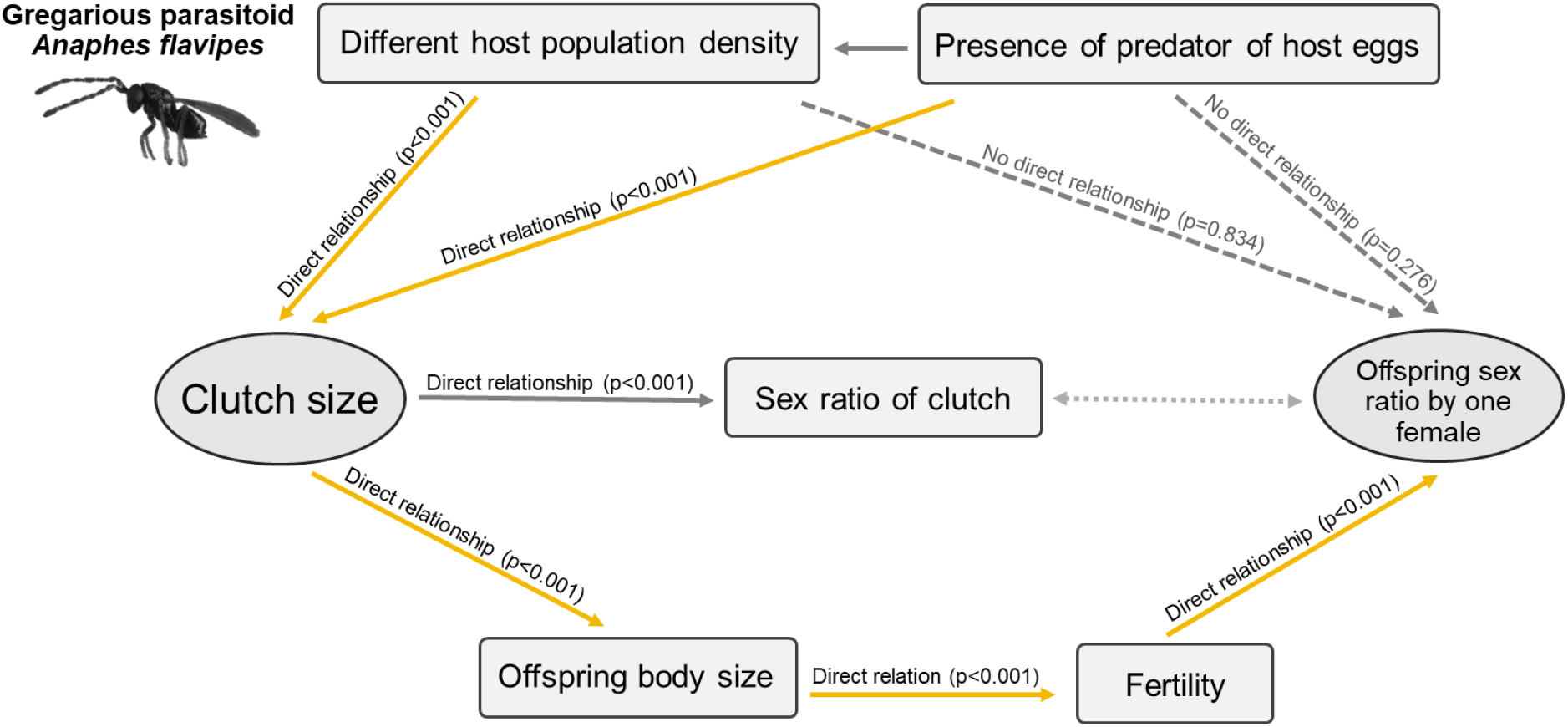
External factors (varying host population density and the presence of the host’s predator) do not directly affect the sex ratio of offspring in the F1 generation, but they do affect the clutch size, which in turn determines the offspring’s body size and their fertility (Samková et al. 2019b). Ultimately, F1’s fertility affects the offspring sex ratio in the F2 generation (yellow lines) - discussed below (Fig. 6).

**Fig. 6.**
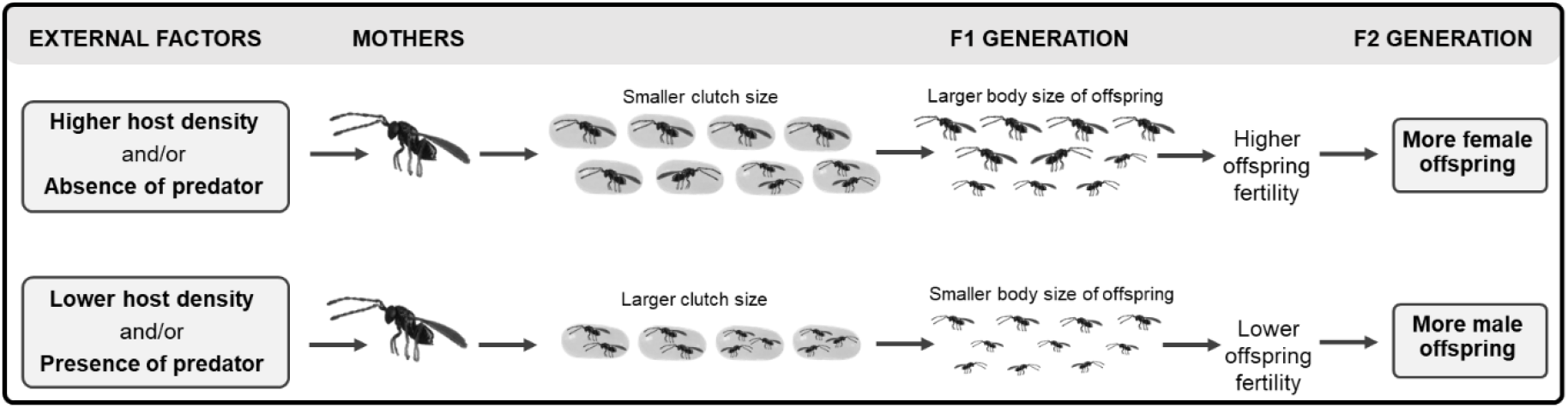
Mothers under different external factors maintain the same offspring sex ratio in the F1 generation (1:4, female 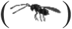, male 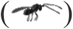), but change the clutch size. The clutch size then determines the offspring’s body size and their fertility (Samková et al. 2019b), and this in turn affects the sex ratio in the F2 generation (this study).

In our previous work, we have shown that the above-mentioned environmental factors alter the clutch size and thus offspring (F1) body size and their fertility (Samková et al. 2019b). Here, we examined the effect of different fertilities of mothers on the offspring sex ratio and showed that with higher fertility, the proportion of females in the offspring increases. The females with a lower fertility seem to choose the safer strategy of having more males in the clutch, ensuring as many mated daughters as possible, since males hatch earlier and mate near the host with females from the same clutch (Anderson & Paschke 1968). Females from a small clutch lacking a male (one individual developing in one host) have to mate randomly with males of other clutches. At the same time, the females from smaller clutches tend to have larger bodies (Bai et al. 1992, Jervis et al. 2008, Boivin & Martel 2012), higher flight efficiency (Visser 1994, Kazmer & Luck 1995), longevity (Banks & Thompson 1985) and longer egg-laying time (Ellers & Jervis 2003) compared to smaller females, all of which increases their probability of mating.

In contrast, in females with a higher fertility, the proportion of female offspring increases, and the likelihood that the female offspring will be mated by a male from the same brood decreases. However, due to the high fertility, we may assume that there will be a high population density of wasps and a higher probability that females will mate randomly with an unrelated male. We have such results for *A. flavipes* in our previous study Samková et al. (2020a, submitted), similar to those in the study of van Alphen & Visser (1990), and King (1993), where the proportion of females offspring increased with superparasitism. Superparasitism simulates a higher population density of wasps (van Alphen & Visser 1990), resulting in a higher probability that unmated females mate randomly.

## CONCLUSION

Our results suggest that parasitoid females prevent overproduction of males and maintain the same sex ratio of their offspring in the F1 generation independently of different clutch sizes caused by changing environmental conditions such as different host population density and the presence of the host’s predator. However, we show the importance of evaluating the fitness of gregarious parasitoids from a two-generation approach. To our previous results, which showed that for gregarious parasitoids with a correlation of clutch size—body size—fertility (i.e., Kazmer & Luck 1995, Wei et al. 2014), the seemingly lower fertility in the F1 generation can eventually lead to a higher fertility in the F2 generation (Samková et al. 2020b), we add here that while the discussed external factors do not affect the offspring sex ratio in the F1 generation, they do have an effect on the F2 generation.

## ACKNOWLEDGEMENTS

We would like to thank the colleagues who lent technical equipment: Petr Šípek, Tomáš Vendl, Zuzana Starostová, Dagmar Říhová, Petr Dolejš, and Pavel Just. We also thank Kateřina Jůzová, Dana Drožová and Bára Křížková for valuable contributions to this work. Finally, we thank Marek Romášek and AJE company for proofreadings the manuscript. This work was supported by the Grant Agency of Charles University (GAUK) (No. 243-227357 to AS), a grant from the Ministry of Education, Youth and Sports of the Czech Republic (no. SVV 260434/2018 to AS and PJ), Charles University Research Centre programme No. 204069 (to PJ), and the Ministry of Agriculture of the Czech Republic, including institutional support MZe-RO0418 and programme NAZV No. QK1910281 (MZe ČR) (to JS).

## REFERENCES

Anderson RC, Paschke JD. 1968. The biology and ecology of *Anaphes flavipes* (Hymenoptera: Mymaridae), an exotic egg parasite of the cereal leaf beetle. Ann Entomol Soc Am. 61:1–5.

Anderson RC, Paschke JD. 1969. Additional Observations on the Biology of *Anaphes flavipes* (Hymenoptera: Mymaridae), with Special Reference to the EfEects of Temperature and Superparasitism on Development. Ann Entomol Soc Am. 62:1316–1321.

Antolin MF. 1992. Sex ratio variation in a parasitic wasp I. Reaction norms. Evolution. 46:1496–1510.

Bai B, Luck RF, Forster L, Stephens B, Janssen JM. 1992. The effect of host size on quality attributes of the egg parasitoid, *Trichogramma pretiosum*. Entomol Exp Appl. 64:37–48.

Banks M, Thomson DJ. 1985. Lifetime mating success in the damselfly *Coenagrion puella*. Anim Behav. 33:1175–1183.

Bates D, Maechler M, Bolker B, Walker S. 2014. lme4: Linear mixed-effects models using Eigen and S4. R package version 1.1–7. URL: http://CRAN.Rproject.org/package=lme4>.

Bezděk J, Baselga A. 2015. Revision of western Palaearctic species of the *Oulema melanopus* group, with description of two new species from Europe (Coleoptera: Chrysomelidae: Criocerinae). Acta Ent Mus Nat Pra. 55:273–304.

Boivin G, Baaren J. 2000. The role of larval aggression and mobility in the transition between solitary and gregarious development in parasitoid wasps. Ecol Lett. 3:469–474.

Boivin G, Martel V. 2012. Size-induced reproductive constraints in an egg parasitoid. J insect physiol. 58:1694–1700.

Carbone S, Nieto MP, Rivera AC. 2007. Maternal size and age affect offspring sex ratio in the solitary egg parasitoid *Anaphes nitens*. Entomol Exp Appl. 125:23–32.

Cloutier C, Duperron J, Tertuliano M, McNeil JN. 2000. Host instar, body size and fitness in the koinobiotic parasitoid *Aphidius nigripes*. Entomol. Exp. Appl. 97:29–40.

Cook JM. 1993. Sex determination in the Hymenoptera: a review of models and evidence. Heredity. 71:421–435.

Dysart RJ, Maltby HL, Brunson, MH. 1973. Larval parasites of *Oulema melanopus* in Europe and their colonization in the United States. Entomophaga. 18:133–167.

Ellers J, Jervis M. 2003. Body size and the timing of egg production in parasitoid wasps. Oikos. 102:164–172.

Farahani HK, Ashouri A, Zibaee A, Abroon P, Alford L. 2016. The effect of host nutritional quality on multiple components of *Trichogramma brassicae* fitness. B Entomol Res. 106:633–641.

Flanagan KE, West SA, Godfray HCJ. 1998. Local mate competition, variable fecundity and information use in a parasitoid. Anim Behav. 56:191–198.

Forsse E, Smith SM, Bourchier RS. 1992. Flight initiation in the egg parasitoid *Trichogramma minutum*: effects of ambient temperature, mates, food, and host eggs. Entomol Exp Appl. 62:147–154.

Godfray HCJ. 1994. Parasitoids: behavioral and evolutionary ecology. Princeton University Press.

Godfray HCJ, Partridge L, Harvey PH. 1991. Clutch size. Annu Rev Ecol Syst. 22:409–429.

Grbić M, Ode PJ, Strand MR. 1992. Sibling rivalry and brood sex ratios in polyembryonic wasps. Nature. 360:254.

Griffiths NT, Godfray HCJ. 1988. Local mate competition, sex ratio and clutch size in bethylid wasps. Behav Ecol Sociobiol. 22:211–217.

Gu H, Wang Q, Dorn S. 2003. Superparasitism in *Cotesia glomerata*: response of hosts and consequences for parasitoids. Ecol Entomol. 28:422–431.

Hamilton WD. 1967. Extraordinary sex ratios. Science. 156:477–488.

Hardy ICW. 1992. Non-binomial sex allocation and brood sex ratio variances in the parasitoid Hymenoptera. Oikos. 65:143–158.

Harrell Jr FE, Harrell Jr MFE. 2019. Package ‘Hmisc’. CRAN R, 235–6.

Harvey JA, Kadash K, Strand MR. 2000. Differences in larval feeding behavior correlate with altered developmental strategies in two parasitic wasps: implications for the size-fitness hypothesis. Oikos. 88:621–629.

Henri DC, van Veen FJF. 2011. Body size, life history and the structure of host-parasitoid networks. Adv Ecol Res. 45:136–180.

Charnov EL, Hartogh RL, Jones WT, van den Assem J. 1981. Sex ratio evolution in a variable environment. Nature. 289: 27–33.

Jervis MA, Ellers J, Harvey JA. 2008. Resource acquisition, allocation, and utilization in parasitoid reproductive strategies. Annu Rev Entomol. 53: 361–385.

Kazmer DJ, Luck RF. 1995. Field tests of the size-fitness hypothesis in the egg parasitoid *Trichogramma Pretiosum*. Ecology. 76:412–425.

King BH. 1989. Host-size-dependent sex ratios among parasitoid wasps: does host growth matter? Oecologia. 78:420–426.

King BH. 1993. Sequence of offspring sex production in the parasitoid wasp, *Nasonia vitripennis*, in response to unparasitized versus parasitized hosts. Anim Behav. 45:1236–1238.

Klomp H, Teerink BJ. 1962. Host selection and number of eggs per oviposition in the egg parasite *Trichgramma embryophagum* Htg. Nature. 195:1020–1021.

Klomp H, Teerink BJ. 1967. The significance of oviposition rates in the egg parasite, *Trichogramma embryophagum* Htg. Arch Neerl Zool. 17:350–375.

Koppik M, Thiel A, Hoffmeister TS. 2014. Adaptive decision making or differential mortality: what causes offspring emergence in a gregarious parasitoid? Entomol Exp Appl 150:208–216.

Kraft TS, van Nouhuys S. 2013. The effect of multi-species host density on superparasitism and sex ratio in a gregarious parasitoid. Ecol Entomol. 38:138–146.

Mawela KV, Kfir R, Krüger K. 2013. Effect of temperature and host species on parasitism, development time and sex ratio of the egg parasitoid *Trichogrammatoidea lutea* Girault (Hymenoptera: Trichogrammatidae). Biol Control. 64:211–216.

Meisner M, Harmon JP, Harvey CT, Ives AR. 2011. Intraguild predation on the parasitoid *Aphidius ervi* by the generalist predator *Harmonia axyridis*: the threat and its avoidance. Entomol Exp Appl. 138:193–201.

Nakashima Y, Birkett MA, Pye BJ, Powell W. 2006. Chemically mediated intraguild predator avoidance by aphid parasitoids: interspecific variability in sensitivity to semiochemical trails of ladybird predators. J Chem Ecol. 32:1989–1998.

Nakashima Y, Senoo N. 2003. Avoidance of ladybird trails by an aphid parasitoid *Aphidius ervi*: active period and effects of prior oviposition experience. Entomol Exp Appl. 109:163–166.

Pandey S, Singh R. 1998. Effect of parental age atcoition on reproduction of *Lysiphlebia mirzai* (Hymenoptera: Braconidae). Entomol gen. 23:187–193.

Pandey S, Singh R. 1999. Host size induced variationin progeny sex ratio of an aphid parasitoid *Lysiphlebia mirzai*. Entomol Exp Appl. 90:61–67.

Polis GA, Myers CA, Holt RD. 1989. The ecology and evolution of intraguild predation: potential competitors that eat each other. Annu Rev Ecol Syst. 20:297–330.

R. Core Team R. A language and environment for statistical computing. R Foundation for Statistical Computing (R Core Team, Vienna, 2019).

Ripley B, Venables B, Bates DM, Hornik K, Gebhardt A, Firth D, Ripley MB. 2013. Package ‘mass’. Cran R, 538.

Rosenheim JA. 1996. An evolutionary argument for egg limitation. Evolution. 50:2089–2094.

Rungrojwanich K, Walter GH. 2000. The Australian fruit fly parasitoid *Diachasmimorpha kraussii* (Fullaway): life history, ovipositional patterns, distribution and hosts (Hymenoptera: Braconidae: Opiinae). Pan-Pacific Entomol. 76:1–11.

Samková A, Hadrava J, Skuhrovec J, Janšta P. 2019a. Reproductive strategy as a major factor determining female body size and fertility of a gregarious parasitoid. J Appl Entomol. 143:441–450.

Samková A, Hadrava J, Skuhrovec J, Janšta P. 2019b. Host population density and presence of predators as key factors influencing the number of gregarious parasitoid *Anaphes flavipes* offspring. Sci Rep-UK. 9:6081.

Samková A, Hadrava J, Skuhrovec J, Janšta P. 2019c. Effect of adult feeding and timing of host exposure on the fertility and longevity of the parasitoid *Anaphes flavipes*. Entomol Exp Appl. 167:932–938.

Samková A, Hadrava J, Skuhrovec J, Janšta P. 2020. Host specificity of the parasitic wasp *Anaphes flavipes* (Hymenoptera: Mymaridae) and a new defence in its hosts (Coleoptera: Chrysomelidae: *Oulema* spp.). Insects. 11:175.

Samková A, Janšta P, Huber TJ. 2017. *Anaphes flavipes* (Foester, 1841) redescription, neotype designation, and comparison with *A. nipponicus* Kuwayama, 1932 (Hymenoptera: Chalcidoidea: Mymaridae). Acta Ent Mus Nat Pra. 57:677–711.

Samková A, Raška J, Hadrava J, Skuhrovec J. 2021a. Scarcity of hosts due to superparasitism indicates an increase of individual offspring fertility by reducing parents’ fertility in gregarious parasitoids. Sci Rep-UK. submitted.

Samková A, Raška J, Hadrava J, Skuhrovec J. 2021b. A new two-generation approach to the population dynamics of gregarious parasitoids. PNAS. submited.

Skuhrovec J, Douda O, Zouhar M, Maňasová M, Nový P, Božik M, Klouček P. 2018. Insecticidal activity of two formulations of essential oils against the cereal leaf beetle. Acta Agr Scand Section B-S P. 68:489–495.

Snyder WE, Ballard SN, Yang S, Clevenger GM, Miller TD, Ahn JJ, Hatten TD, Berryman AA. 2004. Complementary biocontrol of aphids by the ladybird beetle *Harmonia axyridis* and the parasitoid *Aphelinus asychis* on greenhouse roses. Biol Control. 30:229–235.

Schmidt JM, Smith JJB. 1986. Correlations between body angles and substrate curvature in the parasitoid wasp *Trichogramma minutum*: a possible mechanism of host radius measurement. J Exp Biol. 125:271–285.

Stouthamer R, Luck RF, Werren JH. 1992. Genetics of sex determination and the improvement of biological control using parasitoids. Environ Entomol. 21:427–435.

Ueno T. 1999. Host-size-dependent sex ratio in a parasitoid wasp. Researches on Population Ecology. 41:47–57.

Vacari AM, De Bortoli SA, Borba DF, Martins MI. 2012. Quality of *Cotesia flavipes* (Hymenoptera: Braconidae) reared at different host densities and the estimated cost of its commercial production. Biol Control. 63:102–106.

van Alphen JJM, Visser ME. 1990. Superparasitism as an adaptive strategy for insect parasitoids. Annu Rev Entomol. 35:59–79.

Van de Vijver E, Landschoot S, Van Roie M, Temmerman F, Dillen J, De Ceuleners K, Smagghe G, De Baets B, Haesaert G. 2019. Inter-and Intrafield Distribution of Cereal Leaf Beetle Species (Coleoptera: Chrysomelidae) in Belgian Winter Wheat. Environ Entomol. 48:276–283.

Vet LEM, Datema A, Janssen A, Snellen H. 1994. Clutch size in a larval-pupal endoparasitoid: conse-quences for fitness. J Anim Ecol. 63:807–815.

Visser ME. 1994. The importance of being large: the relationship between size and fitness in females of the parasitoid *Aphaereta minuta* (Hymenoptera: Braconidae). J Anim Ecol. 63:963–978.

Waage JK. 1982. Sex ratio and population dynamics of natural enemies-some possible interactions. Ann Appl Biol. 101:159–164.

Waage JK, Lane JA. 1984. The reproductive strategy of a parasitic wasp: II. Sex allocation and local mate competition in *Trichogramma evanescens*. J Anim Ecol. 53:417–426.

Waage JK, Ming NS. 1984. The reproductive strategy of a parasitic wasp: I. optimal progeny and sex allocation in *Trichogramma evanescens*. J Anim Ecol. 53:401–415.

Wei K, Tang YL, Wang XY, Cao LM, Yang ZQ. 2014. The developmental strategies and related profitability of an idiobiont ectoparasitoid *Sclerodermus pupariae* vary with host size. Ecol. Entomol. 39:101–108.

Yu SH, Ryoo MI, Na JH, Choi WI. 2003. Effect of host density on egg dispersion and the sex ratio of progeny of *Bracon hebetor* (Hymenoptera: Braconidae). J Stored Prod Res. 39:385–393.

